# mTORC1 activity oscillates throughout the cell cycle promoting mitotic entry and differentially influencing autophagy induction

**DOI:** 10.1101/2024.02.06.579216

**Authors:** Jay N. Joshi, Ariel D. Lerner, Frank Scallo, Alexandra N. Grumet, Paul Matteson, James H. Millonig, Alexander J. Valvezan

## Abstract

Mechanistic Target of Rapamycin Complex 1 (mTORC1) is a master metabolic regulator that stimulates anabolic cell growth while suppressing catabolic processes such as autophagy. mTORC1 is active in most, if not all, proliferating eukaryotic cells. However, it remains unclear whether and how mTORC1 activity changes from one cell cycle phase to another. Here we tracked mTORC1 activity through the complete cell cycle and uncover oscillations in its activity. We find that mTORC1 activity peaks in S and G2, and is lowest in mitosis and G1. We further demonstrate that multiple mechanisms are involved in controlling this oscillation. The interphase oscillation is mediated through the TSC complex, an upstream negative regulator of mTORC1, but is independent of major known regulatory inputs to the TSC complex, including Akt, Mek/Erk, and CDK4/6 signaling. By contrast, suppression of mTORC1 activity in mitosis does not require the TSC complex, and instead involves CDK1-dependent control of the subcellular localization of mTORC1 itself. Functionally, we find that in addition to its well-established role in promoting progression through G1, mTORC1 also promotes progression through S and G2, and is important for satisfying the Wee1- and Chk1-dependent G2/M checkpoint to allow entry into mitosis. We also find that low mTORC1 activity in G1 sensitizes cells to autophagy induction in response to partial mTORC1 inhibition or reduced nutrient levels. Together these findings demonstrate that mTORC1 is differentially regulated throughout the cell cycle, with important phase-specific functional consequences in proliferating cells.

## Introduction

The cell cycle is a tightly orchestrated series of events that governs cell growth and division. It consists of four phases: G1, S, G2 - which are collectively known as interphase - and mitosis. Each phase has specific biosynthetic requirements that are necessary for successful cell cycle progression, indicating that phase-specific control of cellular metabolism is essential^1^. Cell cycle progression is driven in part by oscillations in the levels of Cyclin proteins, which bind to and activate their respective cyclin-dependent kinases (CDKs)^2^. G1 is a period of low overall CDK activity where cells integrate diverse intracellular and extracellular signals that are required to promote progression through the G1/S restriction point. Once cells enter S phase and begin replicating their DNA, they are committed to dividing and must fulfill all ensuing biosynthetic requirements, making the G1/S transition a critical decision point in the cell cycle^3^. The G1/S transition is facilitated by a sharp increase in Cyclin E levels, which then decrease rapidly again once cells have entered S phase^4^. Subsequently, Cyclin A and Cyclin B levels increase steadily through G2, the cell cycle phase in which cells grow most rapidly^5^. Cyclin B binds and activates CDK1, which is essential for entry into mitosis and for early mitotic events including mitotic spindle formation^6,7^. However, entry into mitosis is restricted by the G2/M checkpoint, which maintains CDK1 in an inactive state through inhibitory phosphorylation of CDK1 by Wee1^8^. The G2/M checkpoint is overcome by activation of the Cdc25 phosphatase which dephosphorylates and activates CDK1, also resulting in Wee1 degradation^9^. Cyclin B levels and CDK1 activity peak early in mitosis, until Cyclin B is targeted for degradation by the anaphase-promoting complex/cyclosome (APC/C), resulting in CDK1 inhibition, completion of mitosis, and return to G1^10,11^.

Mechanistic Target of Rapamycin Complex 1 (mTORC1) is a master metabolic regulator that is active in proliferating eukaryotic cells. Under growth-promoting conditions, mTORC1 coordinates a large-scale anabolic program to synthesize the major macromolecules required for cell growth and proliferation, including proteins, lipids, and nucleic acids, while suppressing catabolic processes such as autophagy^12^. mTORC1 localizes to the cytosolic surface of the lysosome, where it is controlled by two sets of small G proteins, the Rheb and Rag GTPases^13,14^. When Rheb is in its active, GTP-bound state, it directly interacts with and activates mTORC1^15^. Rheb is inhibited by its dedicated GTPase-Activating Protein (GAP), the TSC complex, which also localizes to the lysosomal surface^16^. Hormone and growth factor signals activate mTORC1 by causing the TSC complex to dissociate from the lysosome, thus allowing Rheb activation^16^. The role of the Rag GTPases is to bind and recruit mTORC1 to Rheb at the lysosomal surface^17,18^. The Rags function as nutrient sensors that must be activated by sufficient levels of amino acids, glucose, and cholesterol, to bind and recruit mTORC1^14,19–21^. Thus mTORC1 activity is controlled through spatial regulation of both mTORC1 and the TSC complex in an elegant “AND-gate” mechanism that ensures cells have sufficient nutrients as well as pro-growth signaling inputs to activate mTORC1^13^.

Connections between mTORC1 and the cell cycle have long been proposed, with the mTORC1 inhibitor rapamycin known to cause accumulation of cells in G1^22^. More recently, mTORC1 was reported to be activated downstream of CDK4/6, which are broadly active throughout interphase in complex with Cyclin D^23–25^. Differential regulation of mTORC1 activity has also been reported in mitosis, although with conflicting results, with different studies reporting activation^26^ or inhibition^24,27,28^ of mTORC1 compared to asynchronously growing cell populations. Cell cycle effects on Akt activity have also been reported, although its effects on mTORC1 are unknown^29^. Thus, without a comprehensive characterization of mTORC1 activity through the complete cell cycle, it remains unclear whether and how mTORC1 activity changes from one phase to another.

## Results

### mTORC1 activity oscillates throughout the cell cycle

To determine whether mTORC1 activity changes throughout the cell cycle, we measured mTORC1 activity in each cell cycle phase in HeLa cells. mTORC1 signaling in these cells is dependent on exogenous growth factor, hormone or mitogen signals as demonstrated by suppression of mTORC1 activity upon serum withdrawal (Figure S1A). HeLa cells were synchronized at the G1/S boundary by double thymidine block^30^ and collected at various times after release. Using this synchronization method, we observe an expected increase in cell size as cells progress through S and G2, without a change in cell number (Figure S1B-F). Cells undergo mitosis at 9-10 hrs after release, as observed by the appearance of rounded mitotic cells (Figure S1B). By 14 hrs after release, cell number doubles and cell size is reduced, indicating completion of mitosis and entry into G1 (Figure S1B-F). We further confirmed efficient synchronization and cell cycle progression by quantifying DNA content per cell by flow cytometry for propidium iodide staining (Figure S1C). Greater than 80% of cells are in S-phase at 3 hrs after release, G2 at 7 hrs after release, and G1 at 14 hrs after release (Figure S1C). Synchronized cell cycle progression is also evident by high Cyclin E/low Cyclin B levels at 3 hrs, low Cyclin E/high Cyclin B at 7-9 hrs, and low Cyclin E/low Cyclin B at 14 hrs, corresponding to S, G2, and G1 respectively (Figure 1A).

**Figure 1.**
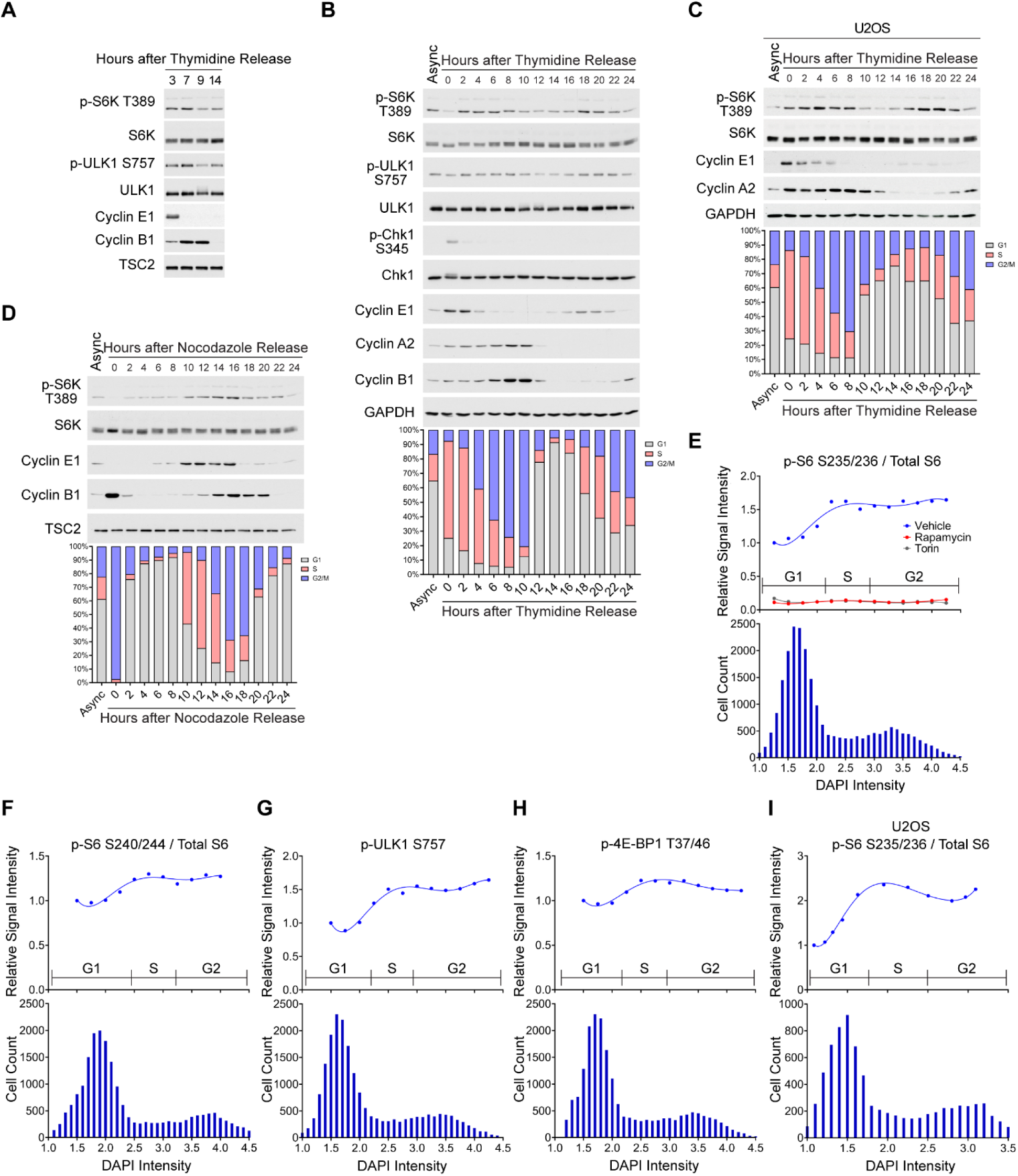
mTORC1 activity oscillates throughout the cell cycle. (A,B) Immunoblots from HeLa cells synchronized at the G1/S boundary by double thymidine block (2mM, 18 hrs each, with 8 hr release in between) and collected at indicated times after release. Cell cycle distribution in (B) was quantified by DAPI staining in parallel samples collected alongside in the same experiment (n=5000-10000 cells per timepoint). (C) U2OS cells were synchronized and collected as in (A,B). (D) HeLa cells were synchronized in mitosis by a single thymidine block (2mM, 18 hr) followed by release into nocodazole (165nM, 12 hr) and then collected at indicated times after release from nocodazole for immunoblotting and cell cycle analysis as in (A). (D) Asynchronous HeLa cells were treated for 2 hrs with vehicle (DMSO), Rapamycin (20 nM), or Torin1 (250 nM) followed by immunofluorescence staining for phospho-S6 S235/236, total S6, and co-staining with DAPI. Staining intensity per cell was quantified (n=24000 cells) and graphed as the average phospho-S6/total S6 within regular DAPI intervals (upper). Cell cycle distribution in the same cells is aligned on the same x-axis scale for reference (lower). (E-G) Asynchronous HeLa cells were immunostained with (E) p-S6 S240/244 and total S6, (F) p-ULK1 S757, (G) p-4EBP1 T37/46 and co-stained with DAPI for analysis as in (D), (n=20000-21000 cells). (I) Asynchronous U2OS cells were immunostained and analyzed as in (D), (n=7900 cells). See also Figure S1.

We measured phosphorylation of the direct mTORC1 substrates S6 Kinase (S6K) and ULK1 and found that they are elevated at the 3 and 7 hr timepoints, corresponding to S and G2 respectively, and lower at 9 and 14 hrs, corresponding to mitosis and G1 respectively (Figure 1A). To further characterize mTORC1 activity through a complete cell cycle, we collected cells every two hours after release from double thymidine block for up to 24 hours, and quantified cell cycle distribution by DNA staining in parallel (Figure 1B). Phospho-S6K and phospho-ULK1 levels are highest from 2 to 8 hrs after release as cells progress through S and G2, then decrease when mitosis occurs by 10 hrs. They remain low as cells progress through G1, before increasing again at 16-18 hours, correlating with the re-induction of Cyclin E1 as cells re-enter the G1/S transition (albeit with some loss of synchronization at these later time points) (Figure 1B). Similar oscillations in mTORC1 activity were observed in U2OS cells and primary human dermal fibroblasts, which also have growth factor-dependent mTORC1 signaling, as well as Ba/F3 murine pro-B cells in which mTORC1 activity is dependent on the cytokine IL-3 (Figure 1C, S1G-M). Despite efficient initial synchronization in S-phase, mTORC1 activity was low in thymidine-arrested cells (Figure 1B, 0 hr time point), likely due to stress from the thymidine block as indicated by phosphorylation of the DNA replication checkpoint protein Chk1, which resolves quickly after release (Figure 1B). As an alternative method, we synchronized cells in mitosis by double thymidine block followed by nocodazole treatment, and again observed that upon release, mTORC1 activity peaks in S/G2 (Figure 1D).

To confirm this effect in unperturbed asynchronous growing cells, we performed immunostaining for phospho-S6 S235/236 and total S6 in HeLa and U2OS cells, as well as phospho-S6 S240/244, phospho-ULK1 S757, and phospho-4E-BP1 T37/46 in HeLa cells, followed by high-throughput quantification of staining intensity per cell with DAPI co-staining to determine cell cycle phase (Figure 1E-I). Each of these mTORC1 activity readouts is elevated to varying degrees in cells in S and G2 compared to G1 (Figure 1E-I). Treatment with the bi-steric mTORC1 inhibitor RMC-5552^31^ in cells synchronized in S, G2, and G1 confirmed that phosphorylation at each of these sites is dependent on mTORC1 in each cell cycle phase (Figure S1N). The cell cycle oscillations in 4E-BP1 and p-S6 S240/244 were not clearly evident by western blot (Figure S1N), consistent with the smaller effect size measured by immunostaining compared to phospho-S6 S235/236 and phospho-ULK1 (Figure 1E-I). Interestingly, in a time course of RMC-5552 treatment, the rate of decrease in phosphorylation of 4E-BP1 and S6 Ser240/244 is slower compared to phospho-S6K, phospho-ULK, and phospho-S6 Ser235/236 (Figure S1N), raising the possibility that slower dephosphorylation kinetics could potentially buffer some mTORC1-mediated signaling events from the cell cycle changes in its activity. Taken together, these data indicate that mTORC1 activity oscillates throughout the cell cycle, peaking in S and G2 and lower in mitosis and G1.

### The TSC complex is required for interphase oscillations in mTORC1 activity

Many signaling inputs to mTORC1 control its activity through effects on the TSC complex. To determine whether the oscillation in mTORC1 activity could be mediated through the TSC complex, we tracked mTORC1 activity throughout the cell cycle in HeLa cells with CRISPR-mediated deletion of the essential TSC complex component TSC2 (sgTSC2). TSC2 protein levels are undetectable in these cells, and mTORC1 activity is growth factor-independent as expected (Figure S2A). sgTSC2 HeLa cells were synchronized at the G1/S boundary by double thymidine block and collected every two hours after release for up to 24 hours (Figure 2A). As in Figure 1, cell cycle distribution was quantified by DNA staining in parallel plates collected alongside in the same experiment. In contrast to control cells (Figure 1), phospho-S6K, phospho-ULK1, and phospho-S6 S235/236 levels do not change throughout interphase in sgTSC2 HeLa cells (Figure 2A). The same result was observed in cells released from synchronization in mitosis by double thymidine block followed by nocodazole, and by quantification of phospho-S6 S235/236 immunostaining in unperturbed asynchronous sgTSC2 HeLa cells (Figure 2B,C). These data indicate that the TSC complex is required for interphase oscillations in mTORC1 activity.

**Figure 2.**
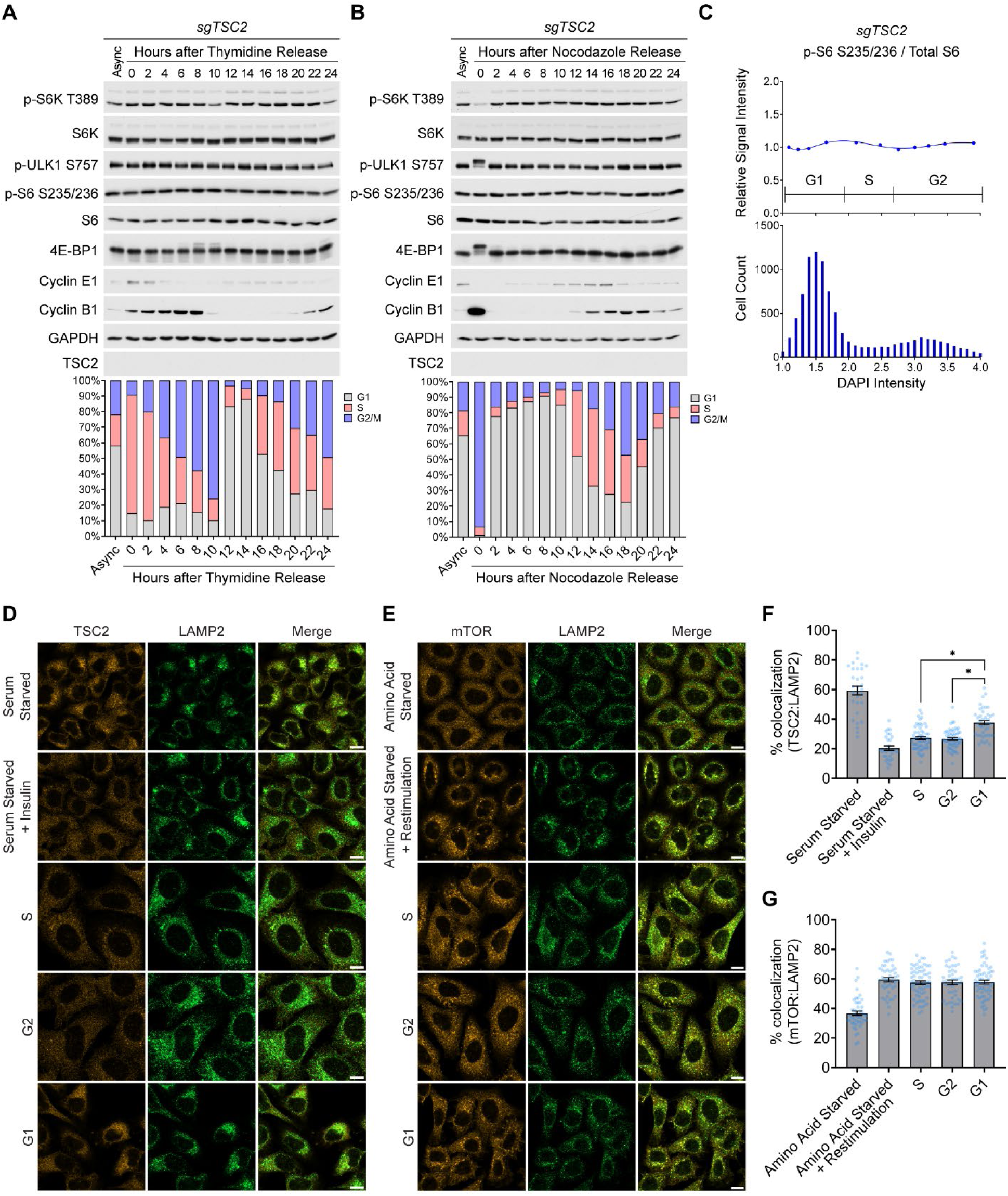
The TSC complex is required for interphase oscillations in mTORC1 activity. (A, B) Immunoblots from HeLa cells with CRISPR-mediated TSC2 deletion (sgTSC2) synchronized at the (A) G1/S boundary by double thymidine block (2mM, 18 hrs each, with 8 hr release in between), or (B) in mitosis by a single thymidine block (2mM, 18 hr) followed by release into nocodazole (165nM, 12 hr), and then collected at indicated times after release. Cell cycle distribution was quantified in parallel samples by (A) DAPI or (B) propidium iodide staining (n=5000-10000 cells per timepoint). (C) Immunofluorescence staining on cells in (A) for phospho-S6 S235/236, total S6, and co-staining with DAPI. Staining intensity per cell was quantified (n=9440 cells) and graphed as the average phospho-S6/total S6 within regular DAPI intervals (upper). Cell cycle distribution in the same cells is aligned on the same x-axis scale for reference (lower). (D) Representative images of HeLa cells serum starved (18 hr) and treated with vehicle or insulin (1 µM, 15 min), or synchronized by double thymidine block and fixed at 3 hrs, 7 hrs and 14 hrs after release for S, G2, and G1, respectively, for co-immunofluorescence staining for TSC2 and LAMP2. (E) Quantification of percent TSC2 colocalized with LAMP2 per cell in (D). Data are presented as mean +/− SEM, n=30-57 cells/group. (F) Representative images of HeLa cells starved of amino acids (50 min), with or without subsequent restimulation (30 min), or synchronized in S, G2, and G1, as in (D), followed by co-immunofluorescence staining for mTOR and LAMP2. (G) Quantification of percent mTOR colocalized with LAMP2 per cell in (F). Data are presented as mean +/− SEM, n=44-63 cells/group. * p < 0.0001 by unpaired Welch’s T-Test. Scalebars, 10 µm. See also Figure S2.

To determine whether lysosomal localization of the TSC complex changes throughout the cell cycle, we performed co-immunofluorescence staining for TSC2 and the lysosomal marker LAMP2 in control HeLa cells synchronized by double thymidine block and collected in S, G2, and G1. Consistent with higher mTORC1 activity in S and G2, TSC2 lysosomal localization is reduced in S and G2 compared to G1 (Figure 2D, 2F). In contrast, mTOR lysosomal localization does not change in S, G2, or G1 (Figure 2E, 2G). Thus the interphase oscillation in mTORC1 activity is mediated through the TSC complex.

### Interphase oscillations in mTORC1 activity are not mediated by Akt, Mek/Erk, or CDK4/6 signaling

Given that the interphase oscillation in mTORC1 activity requires the TSC complex, we examined Akt and Erk signaling, two major inputs that activate mTORC1 through effects on the TSC complex. In HeLa cells synchronized by double thymidine block, Akt activity changes throughout the cell cycle as previously reported^29^, but does not correlate with mTORC1 activity. Based on activating phosphorylation of Akt at Thr308 and Ser473, and phosphorylation of Akt substrates GSK-3 Ser9/21 and TSC2 Thr1462, Akt activity appears slightly higher in G1 when mTORC1 activity is low, (Figure 3A,B, S3A). The Akt inhibitor MK-2206 eliminates Akt phosphorylation and reduces mTORC1 activity in each phase, but mTORC1 activity still remains higher in S and G2 compared to G1, demonstrating that Akt inhibition does not block the oscillation in mTORC1 activity (Figure 3A, p-S6K dark exposure). These data indicate that Akt is a major activator of mTORC1 throughout interphase, but that an Akt-independent mechanism is responsible for the oscillation in mTORC1 activity. The MEK inhibitor trametinib blocks activating phosphorylation of Erk, but does not affect mTORC1 activity in S, G2, or G1, indicating that Erk signaling is not a major input to mTORC1 under these conditions (Figure 3A). Stimulating Akt or Erk signaling, with insulin or EGF, respectively, has similar effects activating Akt and Erk phosphorylation in each phase (Figure 3B). However, mTORC1 activity is still elevated in S and G2 compared to G1, demonstrating that these stimulatory signals do not overcome the cell cycle effect on mTORC1 (Figure 3B). We also tested the role of CDK4/6, which have been previously shown to stimulate mTORC1 activity in studies on asynchronous cell populations^23,24^. Similar to Akt inhibition, CDK4/6 inhibition with abemaciclib reduces mTORC1 activity in each phase, but mTORC1 activity remains higher in S and G2 compared to G1, demonstrating that CDK4/6 inhibition does not block the oscillation in mTORC1 activity (Figure 3C). Amino acid starvation, which inhibits mTORC1 through inactivation of the Rag GTPases^17,18^, fully inhibits mTORC1 in each phase, and amino acid restimulation restores mTORC1 activity back to the respective baseline for each phase (Figure 3D, S3B). These data demonstrate that the cell cycle inputs controlling interphase oscillation in mTORC1 activity are distinct from the above major known upstream mTORC1 regulators.

**Figure 3.**
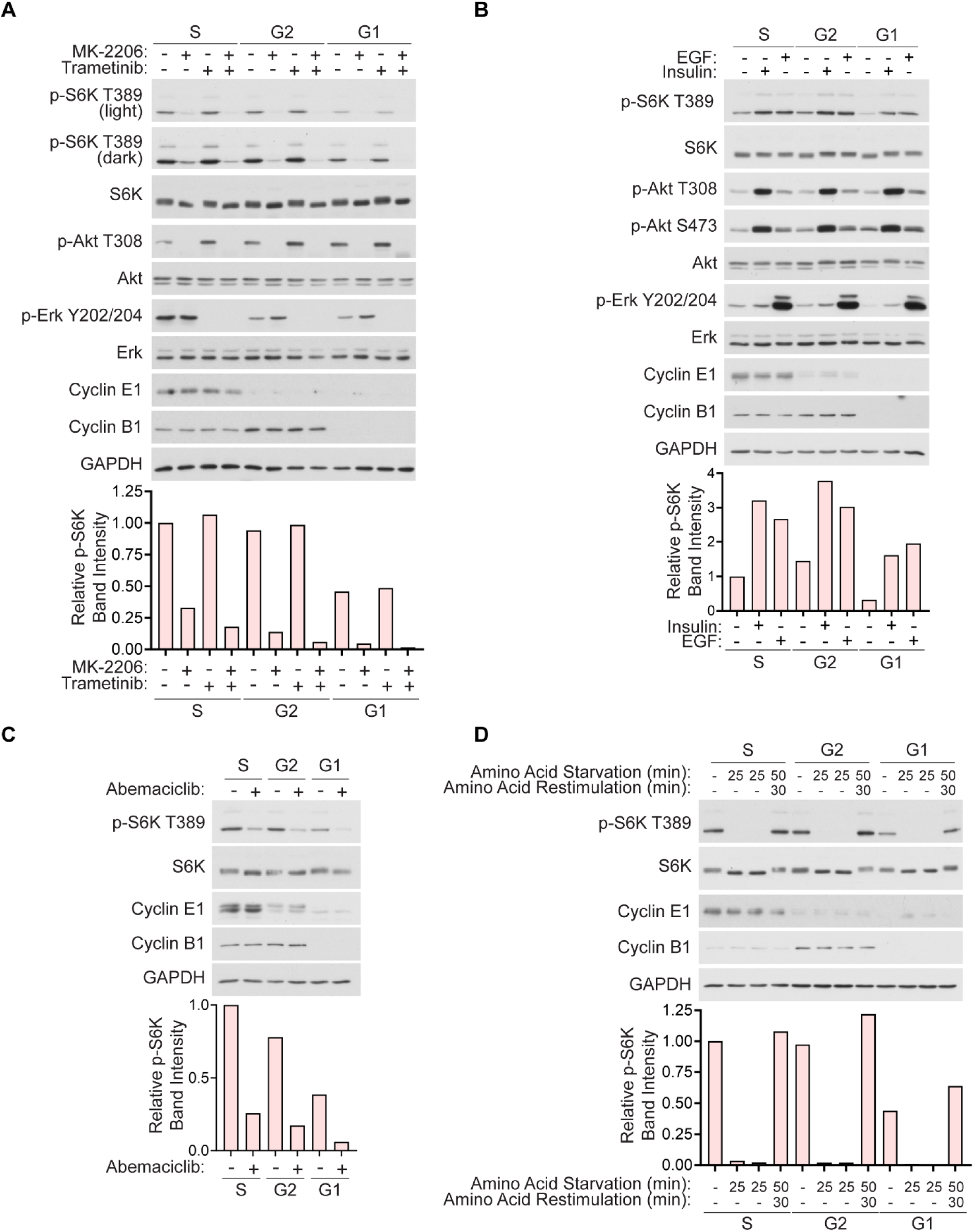
Interphase oscillation in mTORC1 activity is not mediated by Akt or Erk signaling. (A) Immunoblots from HeLa cells synchronized by double thymidine block and treated with the Akt inhibitor MK-2206 (2µM) and/or the MEK inhibitor trametinib (5µM) for 30 min within S, G2, and G1 (2.5 - 3 hrs, 6.5 - 7 hrs, and 13.5 - 14 hrs after release, respectively). (B) Immunoblots from cells synchronized as in (A) and stimulated with insulin (1µM) or EGF (10ng/mL) for 15 min within S, G2, and G1 (2.75 - 3 hrs, 6.75 - 7 hrs, 13.75 - 14 hrs after release, respectively). (C) Immunoblots from cells synchronized as in (A) and treated with the CDK4/6 inhibitor abemaciclib (1µM) for 15 min within S, G2, and G1 as in (B). (D) Immunoblots from cells synchronized as in (A) and starved of amino acids within S, G2, and G1 for indicated times with or without restimulation (beginning at 2 hrs, 6 hrs, and 12 hrs after release, respectively). See also Figure S3.

### mTORC1 activity and lysosomal localization are suppressed in mitosis by CDK1

Although mTORC1 activity does not change throughout interphase in TSC2-deficient cells, S6K phosphorylation is strongly reduced in those cells in mitosis (Figure 2B, 0 hr timepoint). Upon release, S6K phosphorylation increases quickly as cells exit mitosis and enter G1, suggesting mTORC1 activity in mitosis could be suppressed by a TSC complex-independent mechanism. Upward mobility shifts in ULK1 and 4E-BP1 also occur during mitosis (Figure 2B, 0 hr timepoint), which have been previously reported to result from phosphorylation by CDK1^27,32^. Reduced S6K phosphorylation in nocodazole-arrested cells does not appear to be due to nocodazole stress or other indirect effects, as nocodazole treatment in cells that did not receive a prior double thymidine block results in less efficient mitotic synchronization and correspondingly dampened effects on S6K phosphorylation (Figure S2B). Nocodazole treatment in cells that remain arrested in S phase by thymidine block similarly does not strongly affect S6K phosphorylation (Figure S2B). Reduced S6K phosphorylation in mitosis is also evident in TSC2-deficient cells by a small but reproducible decrease in phospho-S6K at the 10 hr timepoint after release from double thymidine block, which is enriched for mitotic cells, as well as a corresponding downward mobility shift in total S6K (Figure 2A).

Given these observations that mTORC1 activity is suppressed in mitosis in TSC2-deficient cells, we measured the lysosomal localization of mTOR during mitosis. HeLa cells were synchronized by double thymidine block and collected in mitosis (9-10 hrs after release), for co-immunofluorescence staining for mTOR, LAMP2, and DAPI. mTOR is dispersed throughout the cell during prophase, metaphase, and anaphase, but re-localizes to lysosomes during telophase/cytokinesis (Figure 4A,B). CDK1 activity peaks early in mitosis but later decreases sharply when the anaphase-promoting complex/cyclosome (APC/C) targets Cyclin B for proteosome-mediated degradation to complete mitosis and cytokinesis^10,11^. As mTOR does not localize to the lysosome early in mitosis, when CDK1 is active, we tested whether CDK1 could suppress mTOR lysosomal localization. Control and sgTSC2 HeLa cells were treated with nocodazole to arrest cells at pro-metaphase, and then treated with the CDK1 inhibitor Ro-3306. Ro-3306 treatment increases lysosomal mTOR localization in both control and sgTSC2 cells (Figure 4C,D), and correspondingly increases mTORC1 activity (Figure 4E). We also confirmed that Ro-3306 treatment in asynchronously growing cells has the expected effects inducing G2/M arrest as previously described^6^ (Figure S4A). CDK1 inhibition does not affect mTORC1 activity or mTOR lysosomal localization in cells in S, G2, or G1 (Figure S4B-D). Taken together these data demonstrate that mTORC1 activity and lysosomal localization are suppressed in mitosis in a CDK1-dependent manner.

**Figure 4.**
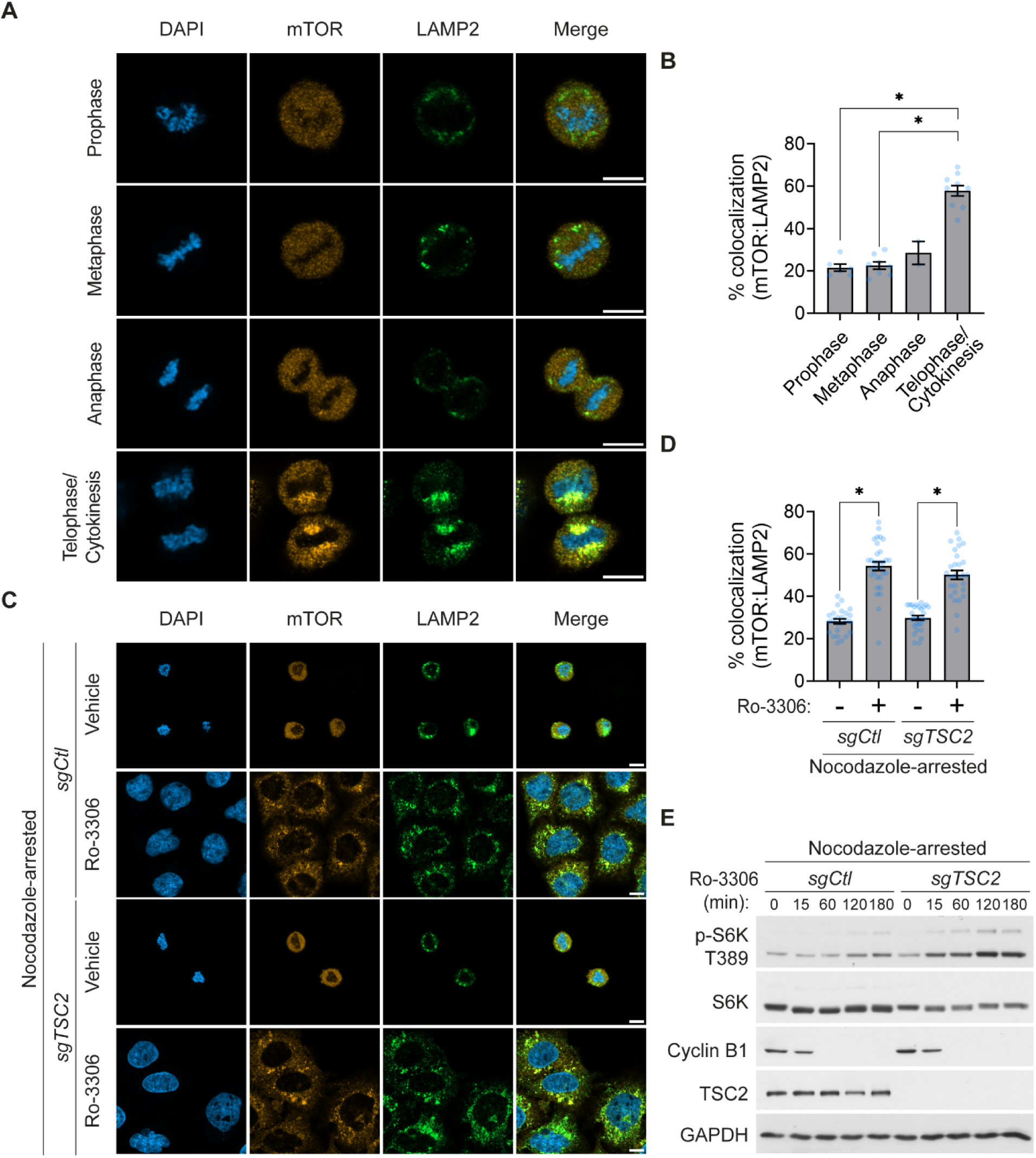
mTORC1 activity and lysosomal localization are suppressed in mitosis by CDK1. (A) Representative images of HeLa cells synchronized by double thymidine block and collected in mitosis for co-immunostaining for mTOR, LAMP2, and DAPI (n=2-10 cells/group). Scalebars, 10 µm. (B) Quantification of percent mTOR colocalized with LAMP2 in (A). (C) Representative images of HeLa cells with control (sgCtl) or CRISPR-mediated TSC2 deletion (sgTSC2) synchronized in mitosis by thymidine block (2mM, 18 hr) followed by release into nocodazole (165nM, 12 hr). Nocodazole-arrested cells were then treated with vehicle (DMSO) or the CDK1 inhibitor Ro-3306 (10µM, 3 hrs) and stained as in (A) (n=25-32 cells/group). Scalebars, 10 µm. (D) Quantification of percent mTOR colocalized with LAMP2 in (C). (E) Immunoblots from control and sgTSC2 HeLa cells synchronized in mitosis as in (C) and treated with vehicle or Ro-3306 (10µM) for indicated times. Graphical data are presented as mean +/− SEM. * p < 0.0001 by unpaired Welch’s T-Test. See also Figure S4.

### mTORC1 promotes progression through S and G2, but not mitosis

mTORC1 is known to promote progression through G1, with extended rapamycin treatment causing cells to accumulate in G1^22^, but whether mTORC1 promotes progression through other cell cycle phases is unknown. Given that mTORC1 activity is elevated in S and G2, we investigated whether mTORC1 can promote progression through those phases as well. We used the bi-steric mTORC1 inhibitor RMC-5552, which potently and specifically inhibits mTORC1, including rapamycin-resistant mTORC1 substrates, without affecting mTORC2 (Figure S1N)^31^. Similar to rapamycin, 24 hr treatment with the bi-steric mTORC1 inhibitor RMC-5552 also causes accumulation of cells in G1 (Figure S5A). We synchronized cells at the G1/S boundary by double thymidine block and measured cell cycle progression after treatment with RMC-5552 in each cell cycle phase. Cells were treated starting 2 hrs after release, when approximately 80% of cells are in S phase, and collected at 5 hrs, when the majority of vehicle-treated cells have progressed to G2 (Figure 5A). RMC-5552 treated cells remained in S phase, indicating that mTORC1 promotes progression through S into G2. Next we began RMC-5552 treatment in early G2 (5 hrs after release) and collected at 9 hrs, when vehicle-treated cells have entered mitosis and G1. In this case RMC-5552 treated cells remained in G2, indicating that mTORC1 also promotes progression through G2 into mitosis (Figure 5B). By contrast, RMC-5552 treatment beginning in late G2/M (9 hrs after release) did not affect progression through mitosis or entry in G1 (Figure 5C). This was confirmed by synchronizing cells in mitosis by double thymidine block followed by nocodazole, where RMC-5552 did not delay completion of mitosis after release (Figure 5D). Similar trends were observed with rapamycin, although to a much lesser extent, consistent with rapamycin only partially inhibiting mTORC1 function (Figure S5B-D). As expected, RMC-5552 and rapamycin treatment in cells synchronized in G1 also blocked progression into S phase (Figure S5E,F). Together these data indicate that in addition to its established role in promoting progression through G1, mTORC1 also promotes progression through S and G2, but is dispensable for progression through mitosis.

**Figure 5.**
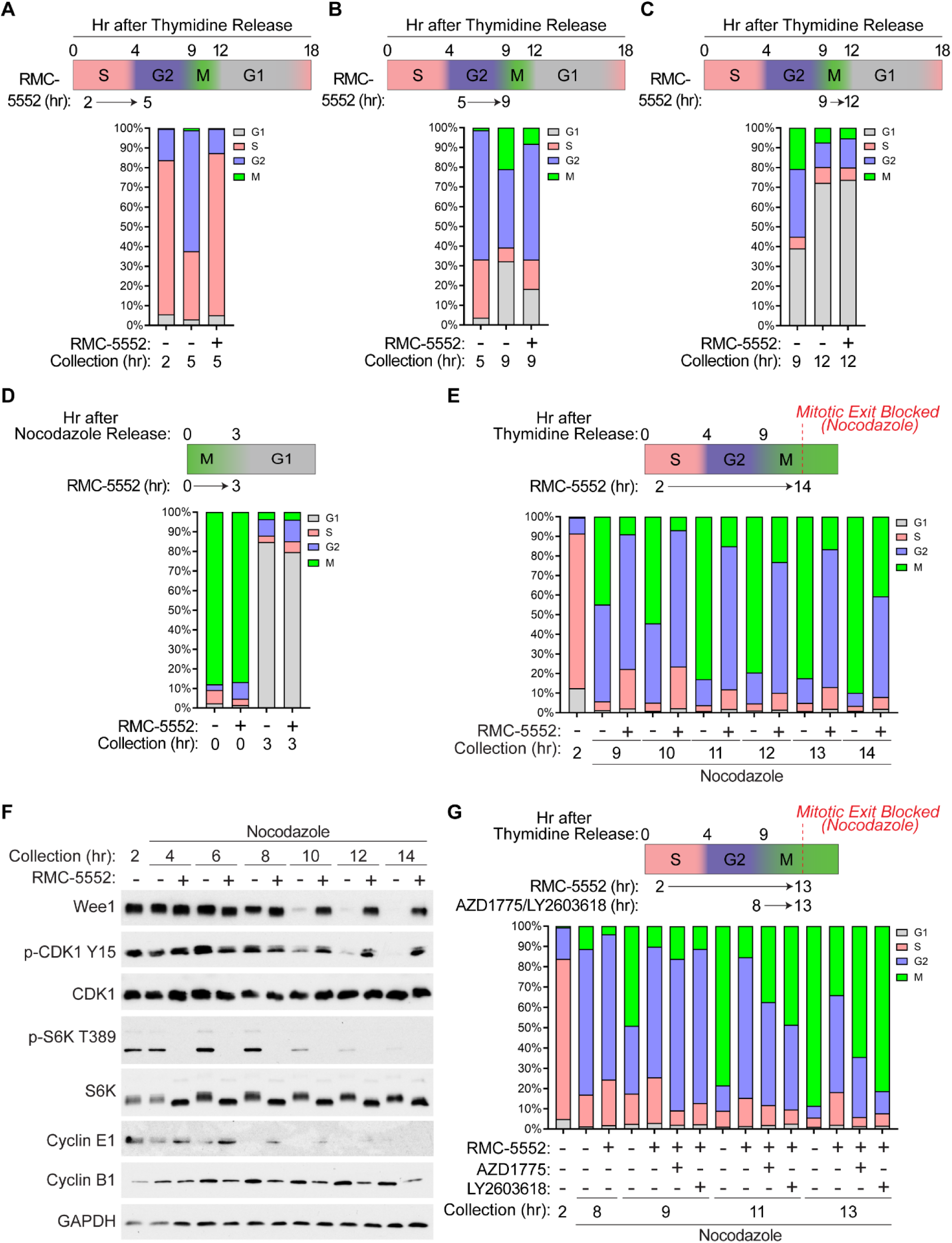
mTORC1 promotes progression through S and G2, but not mitosis. (A-C) HeLa cells were synchronized at the G1/S boundary by double thymidine block and treated with vehicle (DMSO) or the bi-steric mTORC1 inhibitor RMC-5552 (15nM) beginning at (A) 2 hrs, (B) 5 hrs, or (C) 9 hrs after release, and collected at indicated times for quantification of cell cycle distribution by flow cytometry following DAPI staining and p-Histone H3 S10 immunostaining. (D) Cells in (A) were synchronized in mitosis by double thymidine block followed by release into nocodazole (165nM, 12 hr) and treated with vehicle or RMC-5552 (15nM) for 4 hours. Cells were then released from nocodazole in the continued presence of vehicle or RMC-5552 and collected 3 hrs later for quantification of cell cycle distribution as in (A). (E,F) Cells were synchronized as in (A) and treated with vehicle or RMC-5552 (15nM) beginning at 2 hrs after release. Nocodazole (165nM) was added at 2 hrs after release to prevent cells from exiting mitosis. Cells were collected at indicated times for (E) quantification of cell cycle distribution as in (A) or (F) immunoblotting. (G) Cells were synchronized as in (A) and treated with vehicle or RMC-5552 (15nM) beginning at 2 hrs after release, with or without addition of the Wee1 inhibitor AZD1775 (250nM) or the Chk1 inhibitor LY2603618 (500nM) beginning 8 hrs after release. Cells were collected at indicated times for quantification of cell cycle distribution as in (A). Nocodazole (165nM) was also added 2 hrs after release to prevent cells from exiting mitosis. n=5000 cells per condition for flow cytometry experiments. See also Figure S5.

To further characterize the effect of mTORC1 inhibition on cell cycle progression, we synchronized cells at the G1/S boundary by double thymidine block, treated cells with vehicle, RMC-5552, or rapamycin beginning in S phase (2 hrs after release) and collected cells every 2 hrs from 8 to 16 hrs after release. As above, RMC-5552 delayed progression through S and G2, with 80% of vehicle-treated cells having progressed to G1 by 12 hrs after release, compared to only 13% of RMC-5552 treated cells (Figure S5G). RMC-5552 treated cells continue to slowly enter G1 at later time points, suggesting a delay in S and G2 rather than complete arrest.

Rapamycin effects were again less pronounced, with only a slight delay in progression through S and G2 evident at the 8 to 10 hr timepoints (Figure S5G). We performed a similar experiment with the addition of nocodazole to arrest cells once they reach mitosis. 83% of vehicle-treated cells reached mitosis by 11 hrs after release, compared to only 15% of RMC-5552-treated cells (Figure 5E). RMC-5552 treatment beginning in early G2 (5 hrs after release) also delays entry into mitosis, separating the effects in G2 from those in S phase and again demonstrating that mTORC1 promotes progress through both phases (Figure S5H).

The G2/M checkpoint restricts entry into mitosis by regulating the kinase activity of CDK1. Wee1 phosphorylates CDK1 at Tyr15, inactivating CDK1 to prevent entry into mitosis^8,9^. Once the G2/M checkpoint is overcome, phospho-CDK1 levels decrease and Wee1 is degraded^33,34^. We found that RMC-5552 treatment causes CDK1 Tyr15 phosphorylation and Wee1 levels to persist, indicating prolonged activation of the G2/M checkpoint (Figure 5F). Blocking the G2/M checkpoint by inhibiting Wee1 using AZD1775, or by inhibiting Chk1 (which activates Wee1^35–37^) using LY2603618, rescues the delayed mitotic entry caused by RMC-5552 (Figure 5G, S5I).

Immunoblotting in parallel experiments confirmed that AZD1775 has the expected effect blocking CDK1 Tyr15 phosphorylation and that LY2603618 inhibits Chk1 as measured by Chk1 autophosphorylation at Ser296 (Figure S5J,K). Taken together, these data demonstrate that mTORC1 promotes progression through S and G2 and is important for satisfying the G2/M checkpoint to allow entry into mitosis.

### Low mTORC1 activity in G1 sensitizes cells to autophagy induction

mTORC1 phosphorylates and inhibits the autophagy initiating kinase ULK1 to suppress autophagy^38–40^. We investigated whether oscillations in mTORC1 activity influence the induction of autophagy throughout the cell cycle. HeLa cells were synchronized by double thymidine block and treated with rapamycin alone or in combination with bafilomycin A1 for 2 hrs within S, G2, or G1. Bafilomycin A1 blocks autophagosome fusion with lysosomes, allowing accumulation of lipidated LC3 (LC3-II) upon autophagy induction^41,42^. As shown earlier, ULK1 phosphorylation by mTORC1 at Ser757 is lower basally in G1 compared to S and G2 (Figure 6A). Rapamycin partially reduces ULK1 phosphorylation and induces LC3 lipidation in G1 in the presence and absence of bafilomycin A1 (Figure 6A). However, increased LC3 lipidation was not readily detected in S or G2, consistent with higher ULK1 phosphorylation in those phases even in the presence of rapamycin (Figure 6A). Similar results were observed by immunostaining for LC3 puncta (Figure 6B), which are induced during autophagy^43^. We performed the same experiment with RMC-5552, which abolishes ULK1 phosphorylation and thus induces similar LC3 lipidation in S, G2, and G1 (Figure 6C), demonstrating that cells have a similar capacity to induce autophagy in each phase upon complete mTORC1 inhibition. Rapamycin treatment in TSC2-deficient cells, in which mTORC1 activity does not oscillate throughout interphase (Figure 2, 6D, S6A) also similarly induces LC3 lipidation in S, G2, and G1 (Figure 6D), as does RMC-5552 in those cells (Figure S6A). To determine whether differential autophagy induction occurs in response to nutrient deprivation, we switched cells in each phase to a reduced amino acid media (containing 2% of the amino acids present in control media) to reduce, but not completely turn off mTORC1, and observed a similar preferential induction of LC3 lipidation in G1 compared to S and G2 (Figure 6E). Similar results were observed in cells switched to partial EBSS starvation media in each phase (95% EBSS, 5% control media) (Figure S6B). Taken together, these data indicate that low mTORC1 activity in G1, relative to S and G2, sensitizes cells to autophagy induction in response to perturbations that reduce mTORC1 activity.

**Figure 6.**
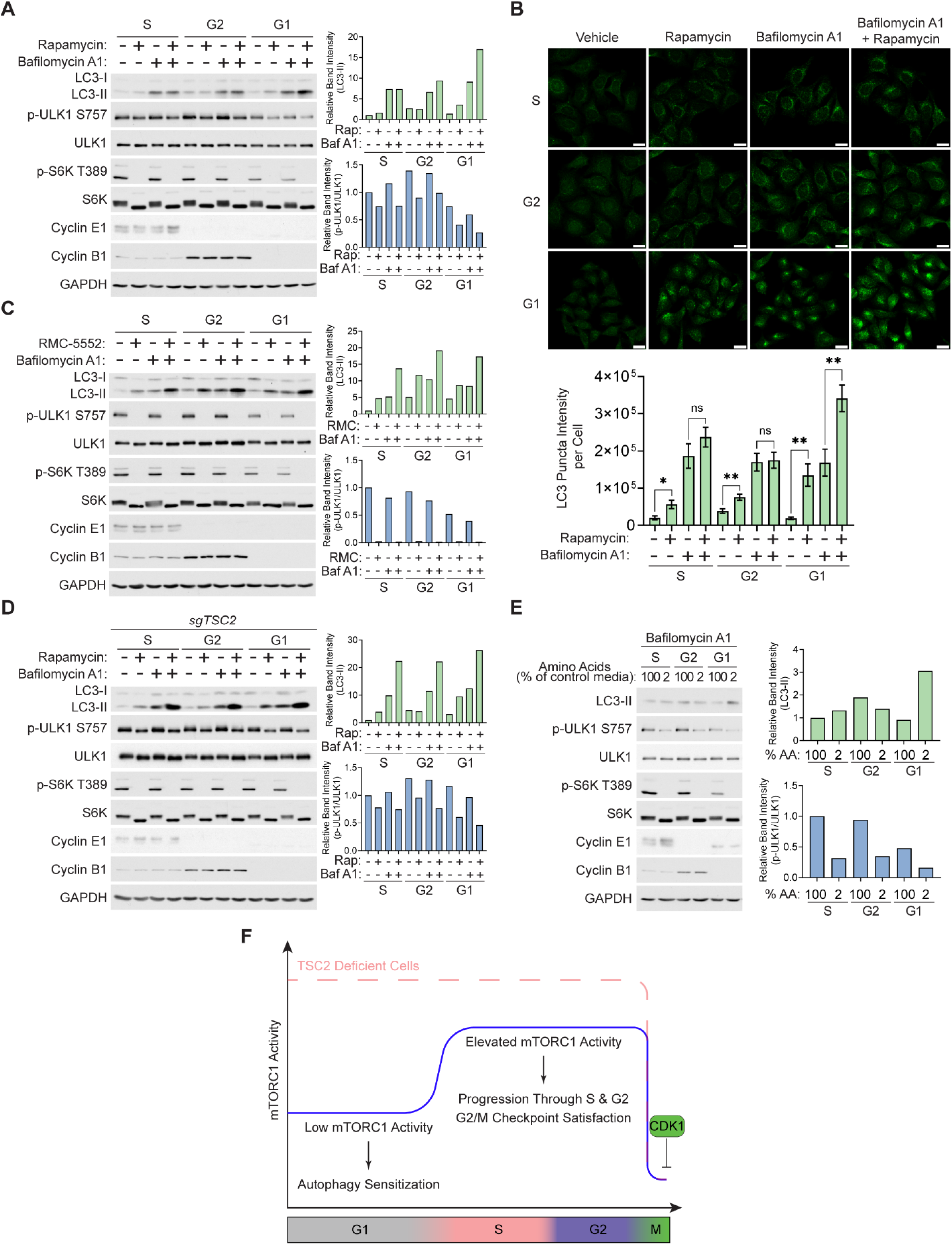
Low mTORC1 activity in G1 sensitizes to autophagy induction. (A) Immunoblots from HeLa cells synchronized by double thymidine block and treated for 2 hr with rapamycin (20nM), in the presence or absence of Bafilomycin A1 (100nM) as indicated, within S, G2, or G1 (1.5 - 3.5 hrs, 5 - 7 hrs, and 13 - 15 hrs after release, respectively). (B) Representative images from HeLa cells synchronized and treated as in (A) followed by immunofluorescence staining for LC3 and quantification of LC3 puncta intensity per cell (n=37-78 cells/group). Scale bars, 20 µM. (C) Immunoblots from HeLa cells synchronized as in (A) and treated with RMC-5552 (15nM), with or without Bafilomycin A1 (100nM) as indicated, within S, G2, or G1 (1.5 - 3.5 hrs, 5 - 7 hrs, and 13 - 15 hrs after release, respectively). (C) HeLa cells with CRISPR-mediated TSC2 deletion synchronized and treated as in (A). (D) HeLa cells synchronized as in (A) and incubated in control or reduced amino acid media (containing 2% amino acids relative to control) for 3 hrs within S, G2 and G1 in the presence of Bafilomycin A1 (100nM). (F) Model. mTORC1 activity oscillates throughout the cell cycle, with low mTORC1 activity in G1 sensitizing to autophagy induction. mTORC1 activity is elevated in S/G2 and promotes progression through those phases. CDK1 represses mTORC1 during mitosis in wild-type and TSC2-deficient cells. Graphical data are presented as mean +/− SEM. * p < 0.001, ** p < 0.0001 by unpaired Welch’s T-Test. See also Figure S6.

## Discussion

Our data demonstrate a model in which mTORC1 activity oscillates throughout the cell cycle, peaking in S and G2, and lowest in mitosis and G1 (Figure 6F). We demonstrate that mTORC1 promotes progression through S and G2, in addition its previously known role in promoting progression through G1. Low mTORC1 activity in G1 also sensitizes cells to autophagy induction (Figure 6F). We found that mTORC1 activity does not change from G1 to S/G2 in TSC2-deficient cells. TSC2 is dissociated from the lysosome in S/G2 compared to G1, consistent with elevated mTORC1 activity in S/G2. Conversely, mTOR lysosomal localization does not change throughout interphase. Together these results suggest that the interphase oscillation in mTORC1 activity is mediated through effects on the TSC complex, rather than changes in nutrient levels or the nutrient sensing machinery that converges to control mTOR localization. Inhibition of Akt, Mek/Erk and CDK4/6 has similar effects on mTORC1 activity in each phase and does not abolish the interphase oscillation mTORC1 activity, indicating that the increase in S/G2 relative to G1 is independent of these known upstream regulators. Thus, these results demonstrate the importance of Akt and CDK4/6 for activating mTORC1 throughout interphase, but suggest an independent cell cycle input further elevates mTORC1 activity in S and G2. This would be predicted to boost anabolic metabolism for the large-scale biosynthesis that occurs during those phases. Conversely, our data also indicate that mTORC1 activity is more dependent on Akt and CDK4/6 in G1 compared to S and G2, as mTORC1 activity is lower upon inhibition of Akt or CDK4/6 in G1 (Figure 3A). The low basal mTORC1 activity in G1 could render cells more sensitive to the upstream signaling inputs that ultimately determine whether cells will progress through the G1/S restriction point. This is supported by Figure 3B where stimulation with insulin or EGF in G1 results in a 5-6-fold increase in S6K phosphorylation, compared to a 2-3-fold increase in S and G2.

While further studies will be necessary to determine the precise mechanism through which mTORC1 activity is elevated in S and G2, we found that the increase in mTORC1 activity correlates with the induction of Cyclin E levels at the G1/S transition. One potential mechanism could involve CDK2 in complex with Cyclin E promoting mTORC1 activation, through either direct or indirect effects on the TSC complex. In this scenario, once Cyclin E levels decrease in S phase, CDK2 might be predicted to sustain mTORC1 activity through G2 by the switch to binding with Cyclin A. It is interesting to note that our high-throughput quantitative immunofluorescence experiments consistently detected a small dip in mTORC1 activity around the S/G2 transition when Cyclin E levels decrease but Cyclin A levels have not yet peaked (Figure 1E,F,G,I). Another non-mutually exclusive possibility is that mTORC1 is partially inhibited in G1 through cell cycle effects that promote activity of the TSC complex.

In contrast to the interphase changes in mTORC1 activity, mTORC1 can be inhibited in mitosis in cells lacking a functional TSC complex (Figure 2B, 4). We find that mTORC1 activity is repressed in mitosis in a CDK1-dependent manner, and that mTOR is dissociated from the lysosome early in mitosis when CDK1 is active, and returns later in mitosis when CDK1 is inactivated (Figure 4A,B). CDK1 inhibition restores mTOR lysosomal localization and increases mTORC1 activity (Figure 4C-E). Interestingly, although mTORC1 activity is similar in control and TSC2-deficient cells arrested in mitosis (Figure 4E, compare 0 hr time points), mTORC1 activity increases more rapidly, and to a greater extent, upon CDK1 inhibition in TSC2-deficient cells (Figure 4E). This indicates that the TSC complex still has a role in repressing mTORC1 during mitosis, and could suggest a dual mechanism of inhibition by CDK1 through both TSC complex-dependent and -independent mechanisms. For reasons that are not entirely clear, CDK1 appears to co-opt parts of the mTORC1 signaling network in mitosis, including directly phosphorylating 4E-BP1 to promote protein synthesis, and ULK1 to maintain suppression of autophagy when mTORC1 activity is low^27,32^. This is in contrast to G1 when CDK1 is inactive, and low mTORC1 activity sensitizes cells to autophagy induction (Figure 6).

Consistent with low mTORC1 activity in mitosis, we also demonstrate that mTORC1 is dispensable for progression through mitosis and entry into G1. Conversely, we find that mTORC1 promotes progression through S and G2, when its activity is elevated (Figure 5). We also find that mTORC1 is important for satisfying the G2/M checkpoint (Figure 5). mTORC1 inhibition prolongs activation of the G2/M checkpoint as indicated by prolonged phosphorylation of Wee1 and CDK1, as well as delayed Wee1 degradation (Figure 5F, S5J,K). Inhibition of Wee1 or Chk1 overcomes the effects of mTORC1 inhibition and promotes entry into mitosis. The G2/M checkpoint ensures that cells do not enter mitosis with damaged or unreplicated DNA^8,9,37^, but has also been implicated as a cell size checkpoint^44,45^, which would be consistent with the role of mTORC1 in stimulating anabolic cell growth. Indeed, mTORC1 inhibition solely in G2 is able to prolong activation of the G2/M checkpoint and delay entry into mitosis (Figure S5H,I,K). These effects are clearly evident using the recently developed bi-steric mTORC1 inhibitor RMC-5552^31^, whereas the allosteric and partial mTORC1 inhibitor rapamycin has relatively minor effects on progression through S and G2 (Figure S5B-D,G), suggesting that rapamycin-resistant functions of mTORC1 are important. Further studies will be necessary to determine the cell cycle phase-specific roles of mTORC1 in controlling cellular metabolism. Given the changing biosynthetic requirements throughout the cell cycle, it is likely that studying metabolic control within specific phases will identify critical new aspects of proliferative metabolism.

## Acknowledgements

We thank Masuda Akther, Ryan Fink, Rita Hahn, Jessica Cervelli, and Dr. Rayees Padder for helpful discussions and technical assistance. We would also like to thank Drs. Brendan Manning and Issam Ben-Sahra for generously sharing reagents. This work was supported by a Department of Defense Tuberous Sclerosis Complex Research Program (TSCRP) Idea Development Award HT9425-23-1-0288 (to A.J.V.), Department of Defense Autism Research Program (ARP) Idea Development Award W81XWH2210806 (to J.H.M.), New Jersey Governor’s Council for Medical Research and Treatment of Autism Award CAUT23BSP014 (to J.H.M.), and an NIH/NCI Cancer Center Support Grant P30CA072720 (provided as a Rutgers Cancer Institute of New Jersey New Investigator Award to A.J.V.).

## Author Contributions

Conceptualization, J.N.J., F.S., and A.J.V.; Methodology, J.N.J, F.S., P.M., and A.J.V.; Validation, J.N.J., A.D.L., F.S., A.N.G., and A.J.V.; Investigation, J.N.J., A.D.L., F.S., A.N.G., and A.J.V.; Resources, J.H.M. and A.J.V.; Writing, J.N.J., J.H.M., and A.J.V.; Visualization, J.N.J. and A.J.V.; Supervision, A.J.V.; Funding Acquisition, A.J.V.

## Methods

### Cell culture

HeLa cells (ATCC #CRM-CCL-2), sgTSC2 HeLa cells (Villa et al., 2021^46^) and U2OS cells (ATCC #HTB-96) were grown in DMEM (Corning #10-017-CV) with 10% heat-inactivated fetal bovine serum (ThermoFisher Scientific #A5256701) and 1% penicillin-streptomycin (Corning #30-002-CI). Primary human dermal fibroblasts (Cell Applications #106-05n) were grown in DMEM with sodium pyruvate (Gibco #10569010), 1% non-essential amino acids (Gibco #11140050), 4µL 2-mercaptoethanol (Gibco #21985023) per 500mL, 10% heat-inactivated fetal bovine serum (ThermoFisher Scientific #A5256701), and 1% penicillin-streptomycin (Corning #30-002-CI). Ba/F3 (DSMZ #ACC-300) cells were grown in RPMI-1640 (Corning #10-040-CV) with 10% heat-inactivated fetal bovine serum (ThermoFisher Scientific #A5256701), 1% penicillin-streptomycin (Corning #30-002-CI), and 10ng/mL IL-3 (R&D systems #403-ML-010).

### Cell cycle synchronization

For double thymidine block, cells were treated with 2 mM thymidine (Sigma #T1895) for 18 hours, then released by washing twice with PBS and incubating in complete media (without thymidine) for 8 hrs. The 2^nd^ thymidine block (2 mM) was then added for another 18 hours before release and collection at time points indicated in the figure legends. For synchronization in mitosis using nocodazole, double thymidine block was performed as above, and then at 5 hours after release from thymidine, 165nM nocodazole (Millipore-Sigma #487928) was added for 12 hours, before release and collection at time points indicated in the figure legends.

### Cell Cycle Distribution

For propidium iodide staining, cells were trypsinized, washed with PBS, then fixed by incubating with pre-chilled (−20°C) 100% ethanol for 15 minutes on ice. Cells were then washed with PBS + 0.1% BSA (Sigma #A7030) and incubated with Propidium Iodide-RNase solution (CST #4087) for 1 hour at room temperature. For DAPI and phospho-Histone H3 (Ser10) staining, cells were fixed in 4% formaldehyde (ThermoFisher Scientific #28908) for 15 min at room temperature, then permeabilized by incubating in pre-chilled (−20°C) 100% methanol for 10 minutes on ice. Cells were then washed with PBS + 0.1% BSA (Sigma #A7030), incubated with anti-phospho-Histone H3 Ser10 primary antibody (CST #9706, 1:50) diluted in PBS + 0.1% BSA for 1 hour at room temperature, washed with PBS again, and then incubated with Alexa Fluor 488-conjugated secondary antibody (ThermoFisher Scientific #A-11001, 1:1000) for 30 minutes at room temperature. Cells were then washed with PBS, incubated with 0.5 µg/mL DAPI (ThermoFisher Scientific #D1306) for 1 minute and then washed again with PBS. Staining intensity was measured using a Gallios Flow Cytometer (Beckman Coulter) and analyzed using Kaluza Analysis 2.1 software.

### Cell counting and cell size measurement

Single cell suspensions were loaded into MOXI Z Type S cassettes (Orflo MXC002-3) and inserted into a Moxi Z Mini Automated Cell Counter (Orflo MXZ001) for measurement of cell volume, cell diameter, and cell counts.

### Chemical compounds

The following compounds were added into the cell culture medium at final concentrations indicated in the figure legends: insulin (Sigma #I9278), EGF (Gibco #PHG0314), rapamycin (Sigma #553210), RMC-5552 (MedChemExpress #HY-132168), abemaciclib (Selleckchem #S7158), trametinib (MedChemExpress #HY-10999), MK-2206 (Selleckchem #S1078), Ro-3306 (APExBIO #A8885), AZD1775 (MedChemExpress #HY-10993), LY2603618 (Selleckchem # S2626), nocodazole (Sigma #1404), thymidine (Sigma #T1895), bafilomycin A1 (MedChemExpress #HY-100558).

### Nutrient starvation

Amino acid free DMEM was made by dissolving amino acid-, glucose-, sodium bicarbonate-, sodium pyruvate-free DMEM powder (US Biological #D9800-28) in water, supplemented with sodium bicarbonate (ThermoFisher Scientific #25080094) to 3.7 g/L concentration, glucose (ThermoFisher Scientific #A2494001) to 4.5 g/L concentration, 10% dialyzed FBS (ThermoFisher Scientific #A3382001), and 1% penicillin-streptomycin (Corning #30-002-CI). pH was adjusted to 7.4. EBSS was purchased from ThermoFisher (#24010-043).

### High-throughput quantitative immunostaining

Cells grown on glass coverslips were fixed with 4% formaldehyde (ThermoFisher Scientific #28908) for 15 min at room temperature, then washed with PBS, and blocked with 5% normal goal serum (CST #5425) plus 0.3% Triton X-100 in PBS for 60 min at room temperature. The following primary antibodies were diluted in PBS with 0.1% BSA (Sigma #A7030) and 0.3% Triton X-100 and added overnight at 4°C: S6 (CST #2317, 1:50), phospho-S6 S235/236 (CST #4858, 1:200), phospho-S6 S240/244 (CST #5364), phospho-ULK1 S757 (CST #14202), and phospho-4E-BP1 T37/46 (CST #2855). Cells were then washed in PBS and incubated with secondary antibodies conjugated to Alexa Fluor 488 (ThermoFisher Scientific #A-11001, 1:1000) and Cy3 (Jackson ImmunoResearch #111-165-144, 1:1000) for 1 hour at room temperature, then stained with 0.5 µg/mL DAPI (ThermoFisher Scientific #D1306). Cells were then washed with PBS and mounted on slides with Fluoromount G (SouthernBiotech #0100-01) for imaging and quantification using the CellInsight CX7 LED Pro High-Content Screening Platform (ThermoFisher Scientific HCSDCX7LEDPRO).

### mTOR and TSC2 Immunofluorescence

Cells grown on glass coverslips were fixed with 4% formaldehyde (ThermoFisher Scientific #28908) for 15 min at room temperature, then washed with PBS. Cells were blocked with Intercept Blocking Buffer (Licor #927-70001) diluted 1:1 in PBS for 60 min at room temperature. Cells were incubated overnight at 4°C in primary antibodies: TSC2 (CST #4308, 1:1250) or mTOR (CST #2938, 1:200) and LAMP2 (Santa Cruz #sc18822, 1:100). Cells were washed with PBS, incubated with secondary antibodies conjugated to Alexa Fluor 488 (ThermoFisher Scientific #A-11001, 1:1000) and Cy3 (Jackson ImmunoResearch #111-165-144, 1:1000) for 1 hour at room temperature, washed with PBS again, and stained with 0.5 µg/mL DAPI (ThermoFisher Scientific #D1306) before mounting on slides with Fluoromount G (SouthernBiotech #0100-01) for imaging using a Zeiss LSM 900 confocal microscope. TSC2:LAMP2 and mTOR:LAMP2 colocalization was quantified using the Zeiss ZenBlue 3.3 analysis suite.

### LC3 Immunofluorescence

Cells grown on glass coverslips were fixed with 100% methanol for 15 min at −20°C, then washed with PBS. Cells were blocked in PBS containing 5% normal goal serum (CST #5425) and 0.3% Triton X-100 for 60 min at room temperature. Cells were incubated overnight at 4°C with LC3 A/B primary antibody (CST #12741, 1:100), then washed with PBS, incubated with Cy3-conjugated secondary antibody (Jackson ImmunoResearch #111-165-144, 1:1000) for 1 hour at room temperature, before washing with PBS again and mounting with Fluoromount G (SouthernBiotech #0100-01). Cells were imaged using a Zeiss LSM 900 confocal microscope. LC3 puncta intensity per cell was quantified using the Zeiss ZenBlue 3.3 analysis suite.

### Immunoblotting

Cells were lysed using lysis buffer containing 20 mM Tris pH 7.5, 140 mM NaCl, 1 mM EDTA, 10% glycerol, 1% Triton X-100, 50 mM NaF, 1 mM DTT, with protease inhibitor cocktail (Sigma #P8340), phosphatase inhibitor cocktail #2 (Sigma #P5726), and #3 (Sigma #P0044) used at 1:100 each. Western blots were performed using the following antibodies purchased from Cell Signaling Technology and used at 1:1000 dilution unless otherwise indicated: phospho-p70 S6 Kinase T389 (CST #9234 1:2000), p70 S6 Kinase (CST #2708, 1:2500), phospho-4E-BP1 T37/46 (CST #2855), 4E-BP1 (CST #9644, 1:5000), phospho-ULK1 S757 (CST #14202), ULK1 (CST #8054), phospho-S6 S235/236 (CST #4858 1:5000), phospho-S6 S240/244 (CST #5364 1:5000), S6 (CST #2317), Cyclin E1 (CST #20808), Cyclin A2 (CST #4656), Cyclin B1 (CST #4138), Cyclin D2 (CST #3741), phospho-TSC2 T1462 (CST #3617), phospho-GSK3 S9/21 (CST #9331), TSC2 (CST #4308), GAPDH (CST #5174, 1:5000), phospho-Akt T308 (CST #13038), phospho-Akt S473 (CST #4060 1:5000), Akt (CST #4691), phospho-Erk Y202/204 (CST #9106), Erk (CST #9102), phospho-Wee1 S642 (CST #4910), Wee1 (CST #13084), phospho-CDK1 Y15 (CST #4539), CDK1 (CST #9116), phospho-Chk1 S345 (CST #2348), phospho-Chk1 S296 (CST #90178), Chk1 (CST #2360), LC3 A/B (CST #12741). Immunoblots were quantified using ImageJ^47^ and normalized to the first lane in each respective blot.

## Quantification And Statistical Analysis

Graphical data are represented as mean ± SEM. p values for pairwise comparisons were determined using an unpaired two-tailed Welch’s t test. Statistical details for individual experiments can be found in their respective figure legends. GraphPad Prism 10 was used to conduct statistical tests and generate graphs.

## Key Resources

| REAGENT or RESOURCE | SOURCE | IDENTIFIER |
| --- | --- | --- |
| **Antibodies** |  |  |
| Phospho-p70 S6 Kinase T389 | Cell Signaling Technology | Cat #9234 |
| p70 S6 Kinase | Cell Signaling Technology | Cat #2708 |
| Phospho-4E-BP1 T37/46 | Cell Signaling Technology | Cat #2855 |
| 4E-BP1 | Cell Signaling Technology | Cat #9644 |
| Phospho-ULK1 S757 | Cell Signaling Technology | Cat #14202 |
| ULK1 | Cell Signaling Technology | Cat #8054 |
| Phospho-S6 S235/236 | Cell Signaling Technology | Cat #4858 |
| Phospho-S6 S240/244 | Cell Signaling Technology | Cat #5364 |
| S6 | Cell Signaling Technology | Cat #2317 |
| Phospho-TSC2 T1462 | Cell Signaling Technology | Cat #3617 |
| TSC2 | Cell Signaling Technology | Cat #4308 |
| Phospho-Akt T308 | Cell Signaling Technology | Cat #13038 |
| Phospho-Akt S473 | Cell Signaling Technology | Cat #4060 |
| Akt | Cell Signaling Technology | Cat #4691 |
| Phospho-Erk Y202/204 | Cell Signaling Technology | Cat #9106 |
| Erk | Cell Signaling Technology | Cat #9102 |
| Phospho-GSK3 S9/21 | Cell Signaling Technology | Cat #9331 |
| Phospho-CDK1 Y15 | Cell Signaling Technology | Cat #4539 |
| CDK1 | Cell Signaling Technology | Cat #9116 |
| Phospho-Wee1 S642 | Cell Signaling Technology | Cat #4910 |
| Wee1 | Cell Signaling Technology | Cat #13084 |
| Phospho-Chk1 S345 | Cell Signaling Technology | Cat #2348 |
| Phospho-Chk1 S296 | Cell Signaling Technology | Cat #90178 |
| Chk1 | Cell Signaling Technology | Cat #2360 |
| GAPDH | Cell Signaling Technology | Cat #5174 |
| LC3 A/B | Cell Signaling Technology | Cat #12741 |
| Cyclin E1 | Cell Signaling Technology | Cat #20808 |
| Cyclin A2 | Cell Signaling Technology | Cat #4656 |
| Cyclin B1 | Cell Signaling Technology | Cat #4138 |
| Cyclin D2 | Cell Signaling Technology | Cat #3741 |
| LAMP2 | Santa Cruz | Cat #sc18822 |
| Alexa Fluor 488-conjugated anti-rabbit secondary antibody | ThermoFisher Scientific | Cat #A-11001 |
| Cy3-conjugated anti-rabbit secondary antibody | Jackson ImmunoResearch | Cat #111-165-144 |
| **Chemicals and recombinant proteins** |  |  |
| Thymidine | Sigma | Cat #T1895 |
| Nocodazole | Sigma | Cat # M1404 |
| DAPI | ThermoFisher Scientific | Cat #D1306 |
| Rapamycin | Sigma | Cat #553210 |
| RMC-5552 | MedChemExpress | Cat #HY-132168 |
| MK-2206 | Selleckchem | Cat #S1078 |
| Trametinib | MedChemExpress | Cat #HY-10999 |
| Abemaciclib | Selleckchem | Cat #S7158 |
| AZD1775 | MedChemExpress | Cat #HY-10993 |
| LY2603618 | Selleckchem | Cat #S2626 |
| Bafilomycin A1 | MedChemExpress | Cat #HY-100558 |
| Ro-3306 | APExBIO | Cat #A8885 |
| EGF | Gibco | Cat #PHG0314 |
| IL-3 | Sigma | Cat #I9278 |
| **Experimental models: Cell lines** |  |  |
| HeLa | ATCC | Cat #CRM-CCL-2 |
| sgCtl HeLa | Provided by Issam Ben-Sahra Lab | Villa et al., 2021<sup>46</sup> |
| sgTSC2 HeLa | Provided by Issam Ben-Sahra Lab | Villa et al., 2021<sup>46</sup> |
| U2OS | ATCC | Cat #HTB-96 |
| Ba/F3 | DSMZ | Cat #ACC-300 |
| Primary human dermal fibroblasts | Cell Applications | Cat #106-05n |
| **Software** |  |  |
| ImageJ | National Institutes of Health<sup>47</sup> | https://imagej.nih.gov/ij/ |
| ZenBlue 3.3 | Zeiss | https://www.zeiss.com/microscopy/en/products/software/zeiss-zen.html |
| HCS Studio 5.0 Cell Analysis | ThermoFisher Scientific |  |
| Prism 10.0 | GraphPad | https://www.graphpad.com/ |
| Kaluza 2.1 | Backman Coulter | https://www.beckman.com/flow-cytometry/software/kaluza |
| **Other** |  |  |
| EBSS | ThermoFisher Scientific | Cat #24010-043 |
| Normal Goat Serum | Normal Goat Serum | Cat #5425 |
| DMEM | Corning | Cat #10-017-CV |
| Fetal Bovine Serum | ThermoFisher Scientific | Cat #A5256701 |
| Intercept Blocking Buffer | Licor | Cat #927-70001 |

## Supplemental Figure Legends

**Figure S1.**
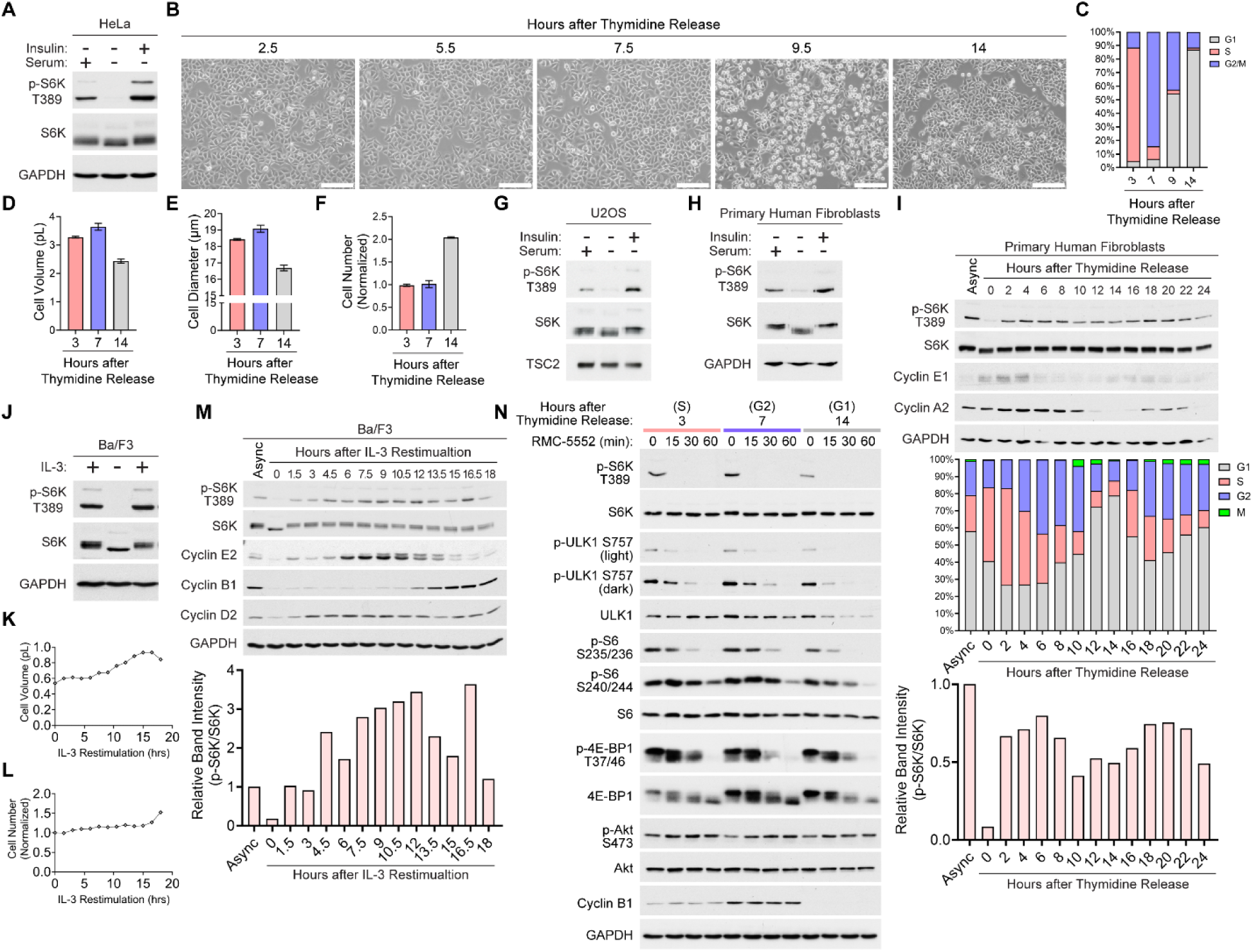
Supplemental data supporting Figure 1. (A) Immunoblots from HeLa cells serum starved for 18 hours then treated with insulin (1 µM, 15 min). (B) Representative images of HeLa cells synchronized by double thymidine block and released for indicated times. Scalebars, 0.5 mm. (C) HeLa cells were synchronized as in (B), and cell cycle distribution was quantified by flow cytometry on propidium iodide-stained cells. (D-F) HeLa cells were synchronized as in (B) and collected at indicated times for quantification of (D) cell volume, (E) cell diameter and (F) viable cell counts. (G,H) Immunoblots from (G) U2OS cells and (H) primary human fibroblasts treated as in (A). (I) Immunoblots from primary human fibroblasts synchronized by double thymidine block and collected at indicated times after release. Cell cycle distribution was quantified by flow cytometry following DAPI staining and phospho-Histone H3 Ser10 immunostaining in parallel samples collected alongside in the same experiment (n=3000-5000 cells per timepoint). (J) Ba/F3 cells were starved of IL-3 for 22 hrs and then acutely restimulated with IL-3 (10ng/mL, 15 min). (K-M) Ba/F3 cells were arrested in G1 by IL-3 deprivation and then collected at indicated times following IL-3 restimulation for (K) cell volume measurement, (L) viable cell counting, and (M) immunoblotting. (N) Immunoblots from HeLa cells synchronized as in (B) and collected at indicated times after release, following treatment with RMC-5552 (15nM) for indicated times.

**Figure S2.**
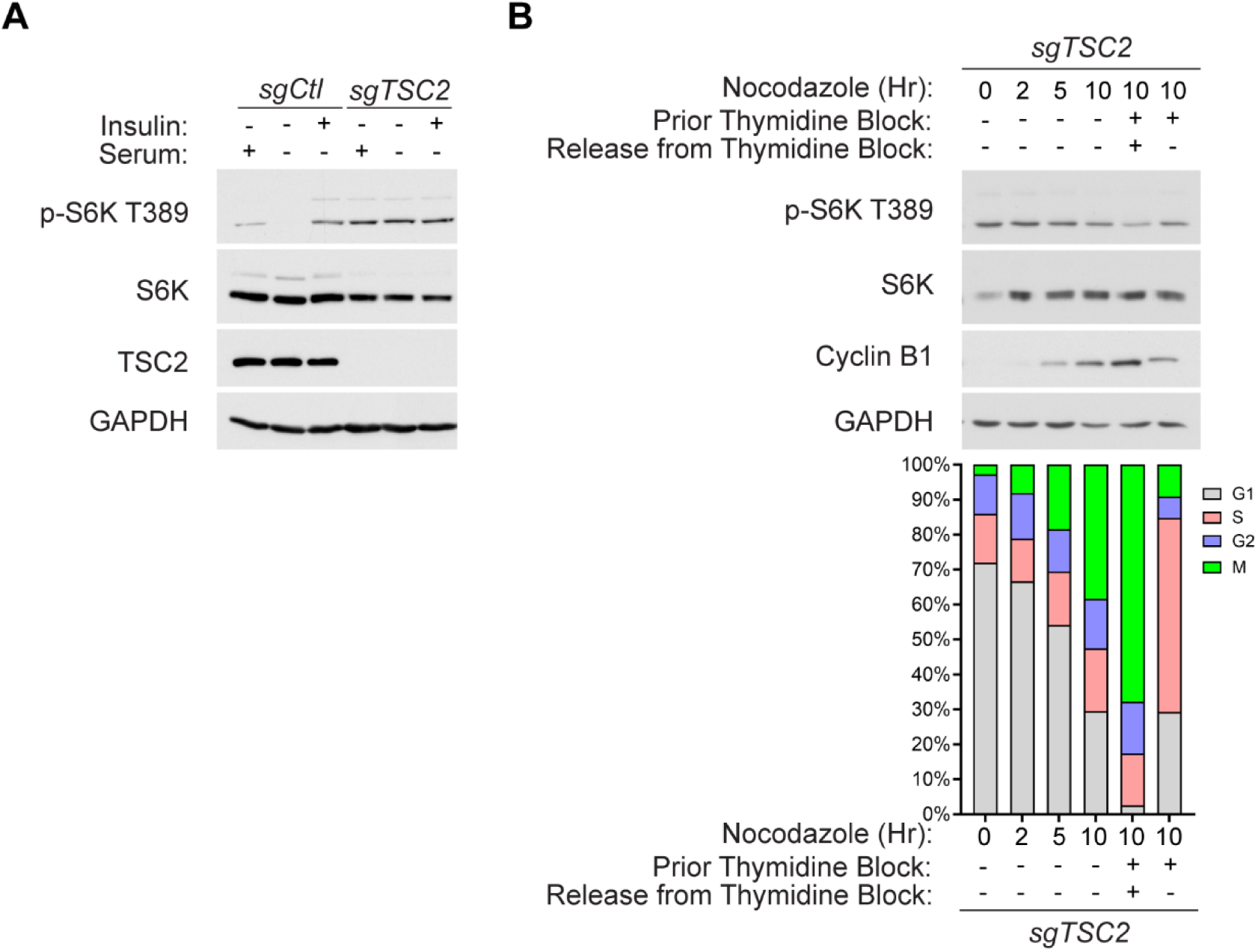
Supplemental data supporting Figure 2. (A) Immunoblots from HeLa cells with control (sgCtl) or CRISPR-mediated TSC2 deletion (sgTSC2) serum starved for 18 hours then treated with insulin (1µM, 15 min). (B) sgTSC2 HeLa cells were treated with nocodazole (165nM) for indicated times (Lanes 1-4). In Lane 5 cells were treated with a single thymidine block (2mM, 18 hr) followed by release into nocodazole (165nM, 12 hr) to synchronize cells in mitosis. Cells in Lane 6 were treated as in Lane 5, except without release from thymidine so cells could not progress out of S phase. Cell cycle distribution was quantified in parallel samples collected alongside in the same experiment by flow cytometry for propidium iodide and phospho-Histone H3 Ser10 immunostaining to identify mitotic cells (n=5000 cells per condition).

**Figure S3.**
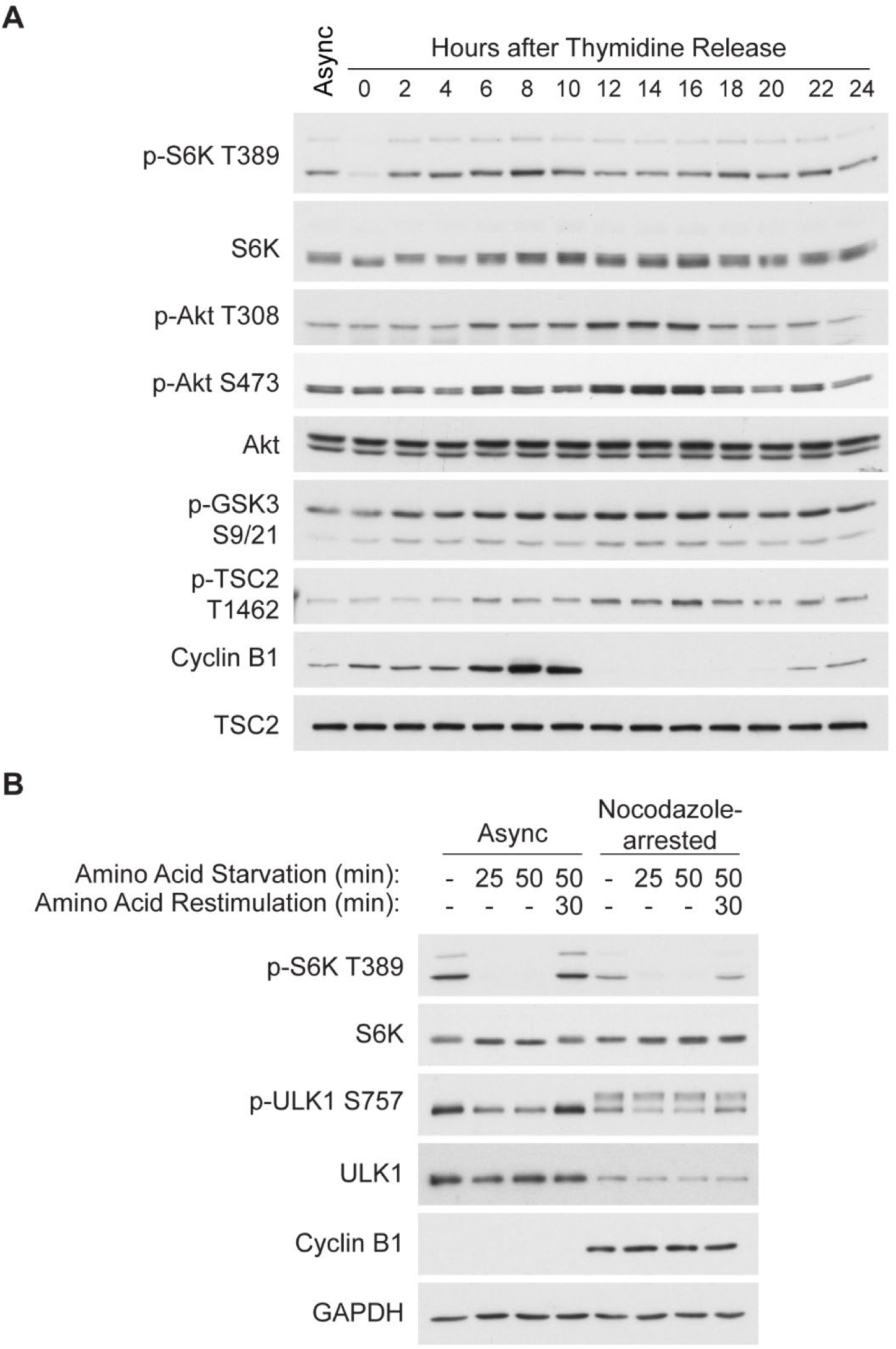
Supplemental data supporting Figure 3. (A) Immunoblots from HeLa cells synchronized at the G1/S boundary by double thymidine block (2mM, 18 hrs each, with 8 hr release in between) and collected at indicated times after release. (B) Immunoblots from asynchronous HeLa cells and cells synchronized in mitosis by a single thymidine block (2mM, 18 hr) followed by release into nocodazole (165nM, 12 hr), starved of amino acids for indicated times with or without restimulation.

**Figure S4.**
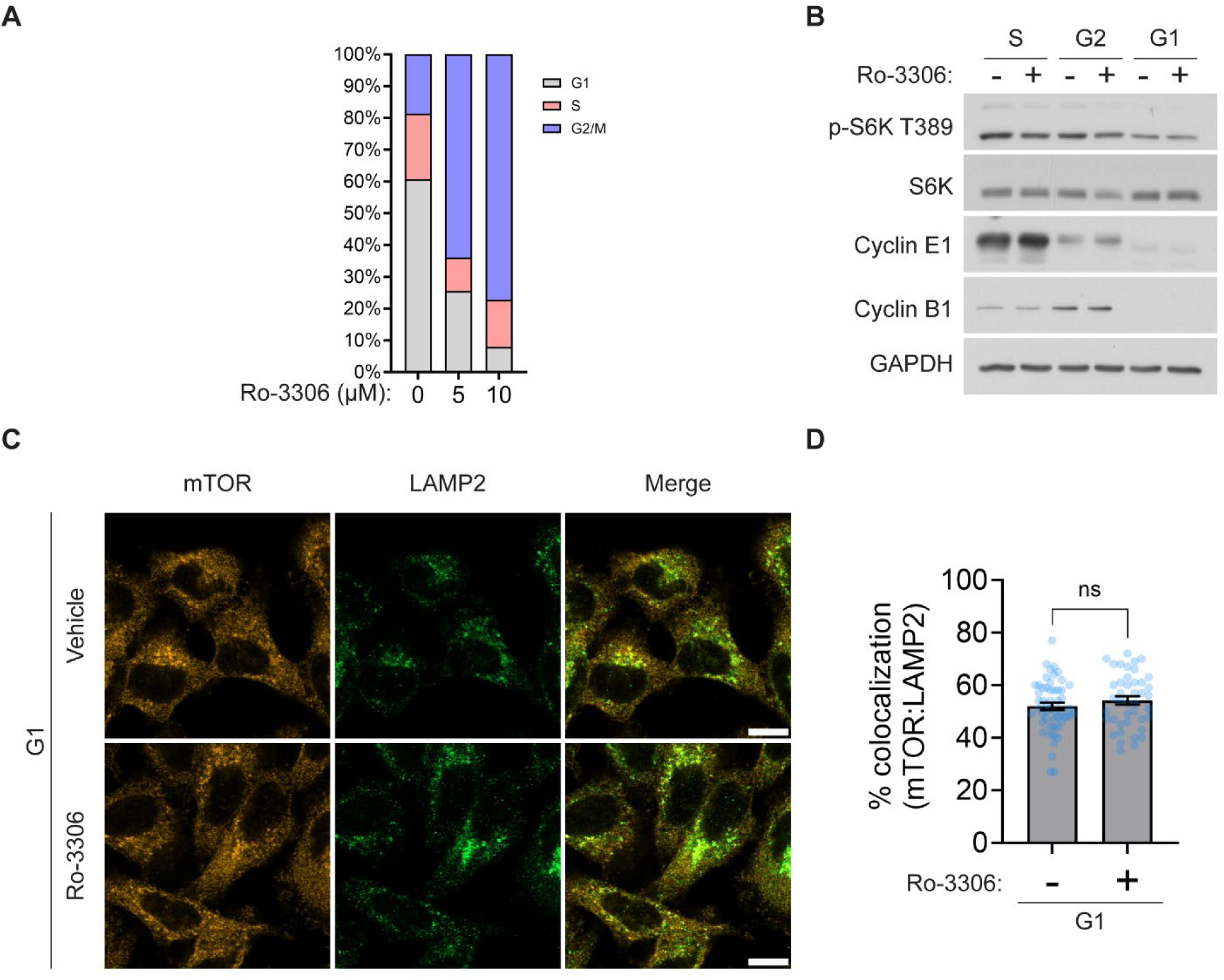
Supplemental data supporting Figure 4. (A) HeLa cells were treated with vehicle (DMSO) or the CDK1 inhibitor Ro-3306 for 18 hrs at indicated concentrations. Cell cycle distribution was quantified by flow cytometry on propidium iodide-stained cells. (B) Immunoblots from HeLa cells synchronized by double thymidine block and treated with Ro-3306 (10µM) for 2 hrs within S, G2, and G1 (1.5 - 3.5 hrs, 5 - 7 hrs, and 12 - 14 hrs after release, respectively). (C) Representative images of HeLa cells synchronized by double thymidine block and treated with Ro-3306 (10µM) for 3 hrs within G1 (12 - 15 hrs after release) for co-immunofluorescence staining for mTOR and LAMP2 (n=42-52 cells/group). Scalebars, 10 µm. (D) Quantification of percent mTOR colocalized with LAMP2 in (C). Data are presented as mean +/− SEM. ns=not significant by unpaired Welch’s T-Test.

**Figure S5.**
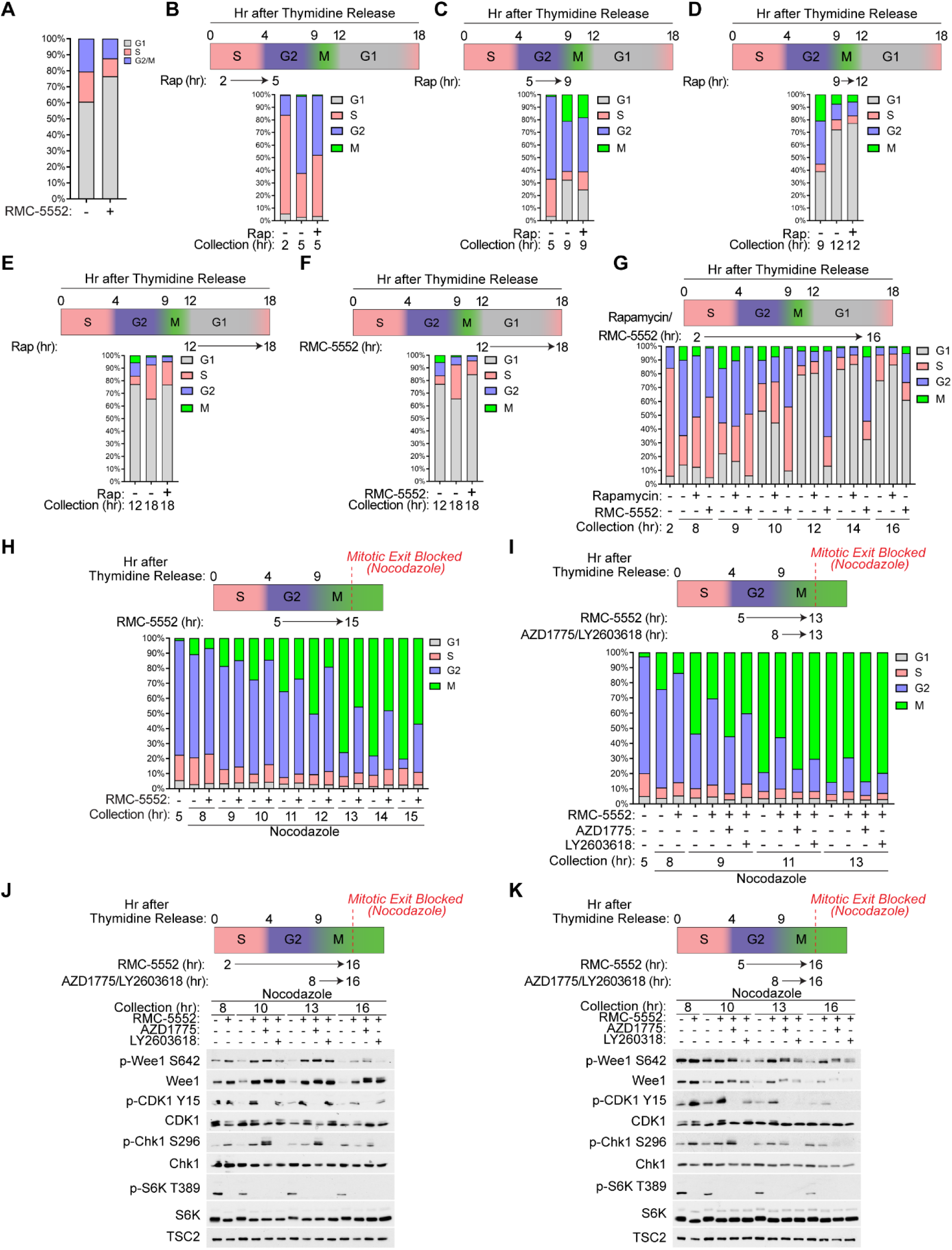
Supplemental data supporting Figure 5. (A) HeLa cells treated with vehicle or RMC-5552 (15nM) for 24 hours. Cell cycle distribution was quantified by flow cytometry following propidium iodide staining. (B-E) HeLa cells were synchronized at the G1/S boundary by double thymidine block and treated with vehicle (DMSO) or rapamycin (20nM) beginning at (B) 2 hrs, (C) 5 hrs, (D) 9 hrs, or (E) 12 hrs after release, and collected at indicated times for quantification of cell cycle distribution by flow cytometry following DAPI staining and p-Histone H3 Ser10 immunostaining. Vehicle treated samples in Figure S5B-D are the same as in Figure 5A-C, respectively, which were done together in the same experiment. (F) HeLa cells were synchronized as in (B) and treated with vehicle or RMC-5552 (15nM) from 12-18 hr after release for quantification of cell cycle distribution as in (B). (G) HeLa cells were synchronized as in (C) and treated with vehicle, rapamycin (20nM) or RMC-5552 (15nM) beginning 2 hrs after release and collected at indicated times for quantification of cell cycle distribution as in (C). (H) Cells were synchronized as in (B) and treated with vehicle or RMC-5552 (15nM) beginning at 5 hrs after release for collection at indicated times. Nocodazole (165nM) was added at 5 hrs after release to prevent cells from exiting mitosis. Cell cycle distribution was quantified as in (B). (I) Cells were synchronized as in (B) and treated with vehicle or RMC-5552 (15nM) beginning at 5 hrs after release, with or without addition of the Wee1 inhibitor AZD1775 (250nM) or the Chk1 inhibitor LY2603618 (500nM) beginning at 8 hrs after release. Cells were collected at indicated times for quantification of cell cycle distribution as in (B). Nocodazole (165nM) was added 5 hrs after release to prevent cells from exiting mitosis. (J,K) Immunoblots from cells synchronized as in (B) and treated with vehicle or RMC-5552 (15nM) beginning at (J) 2 hrs after release or (K) 5 hrs after release, with or without addition of the Wee1 inhibitor AZD1775 (250nM) or the Chk1 inhibitor LY2603618 (500nM) beginning at 8 hrs after release. Cells were collected at indicated times. Nocodazole (165nM) was added 5 hrs after release to prevent cells from exiting mitosis. n=5000 cells per condition for flow cytometry experiments.

**Figure S6.**
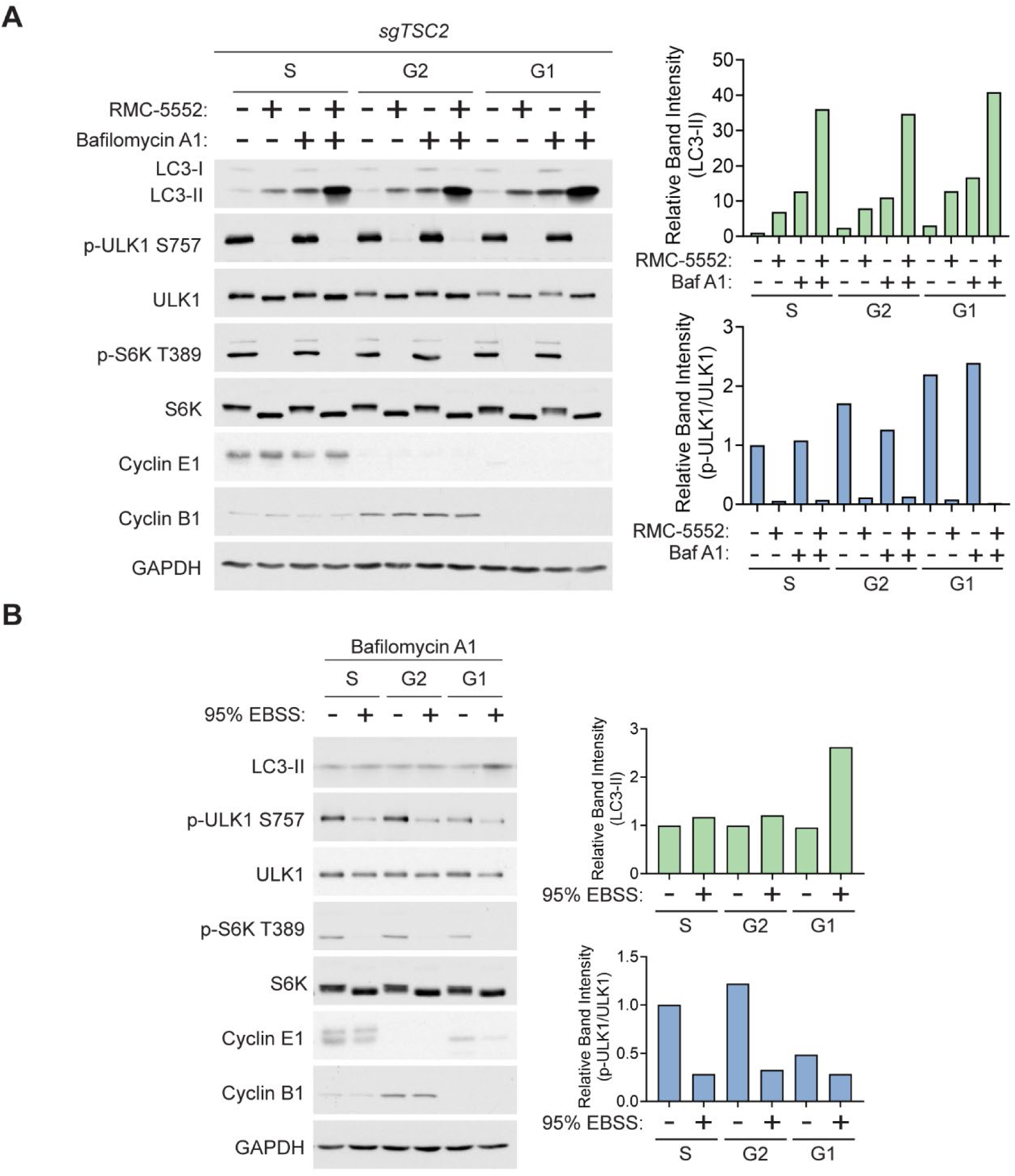
Supplemental data supporting Figure 6. (A) Immunoblots from HeLa cells with CRISPR-mediated TSC2 deletion synchronized by double thymidine block and treated for 2 hr with RMC-5552 (15nM), with or without Bafilomycin A1 (100nM) as indicated, within S, G2, or G1 (1.5 - 3.5 hrs, 5 - 7 hrs, and 13 - 15 hrs after release, respectively). (B) HeLa cells synchronized as in (A) and incubated in control (-) or 95% EBSS starvation media with 5% control media (+) for 3 hr within S, G2 and G1 in the presence of Bafilomycin A1 (100nM).

